# Shotgun proteomic analysis of the caterpillar *Lonomia obliqua* (Lepitoptera, Saturniidae) hemolymph and effects in rat hippocampal neurons culture

**DOI:** 10.1101/2023.03.20.533482

**Authors:** Silviane Maggi, Antônio Frederico Michel Pinto, Mariana Sayuri Berto Udo, Mariana Aguilera Alencar da Silva, Raphael Caio Tamborelli Garcia, Luciane Minetto, Leandro Tasso, Pablo Machado, Diógenes Santiago Santos, Thiago Barcellos, Pedro Ismael da Silva Junior, Ronaldo Zucatelli Mendonça, Tania Marcourakis, Sidnei Moura

## Abstract

Study of substances with potentially neuroprotective has been one of the research focus on drugs development. Toxic proteins of *Lonomia obliqua* caterpillars, which have caused several accidents in southern Brazil, were identified in the hemolymph with anti-apoptotic activity. This study aims the evaluation of the protein profile and the hemolymph effect on cell viability of rats’ primary cultured hippocampal neurons after apoptosis induction. Semi-quantitative shotgun proteomics approach was used to evaluate the protein profile of 3 caterpillars lots of different origin. Were identified a total of 76 proteins, 71 in hemolymph and 40 in fractions. Antiviral protein predominated in crude hemolymph, following by serine proteases, hemolins and protease inhibitors. In fractions were identified hemolins, serine proteases and protease inhibitors. The treatment of rats’ primary cultured hippocampal neurons with the chromatographic fraction at concentration of 0.05 and 0.10% (v/v) for 24 hours, with subsequently apoptosis induction was able to maintain cell viability significantly higher than positive control. Hemolymph protein composition can show qualitative and quantitative variations intra species when compared different origins animals and consequently exposed to various environmental factors. The results shown on this study may contribute to the identification of proteins with potential use as neuroprotective in degenerative conditions.

## 1. Introduction

Insects represent about 75% of all invertebrate animals, are among the more adapted to life on earth. These organisms are capable of producing a number of chemicals that help them survive the environmental attacks during their evolution (Ratcliffe *et al.*, 2011). Thus, evolutionarily, poisons and venoms animals were developed for the purpose of defense against predators or prey capture, causing physiological changes in natural enemies. However, they have potential application as new therapeutic drugs and have been the subject of research (Calvete, 2009).

Several accidents involving the larval form of the *Lonomia obliqua* Walker, 1855 (Lepidoptera, Saturniidae) have been reported in southern Brazil, since the 80s (Duarte *et al.*, 1990; Duarte *et al.*, 1994; Kelen *et al.*, 1995; Duarte *et al.*, 1996; Zannin *et al.*, 2003). Individuals who had contact with bristles of *L. obliqua* caterpillars can manifest a hemorrhagic syndrome associated with consumption coagulopathy which may include intravascular hemolysis and acute renal failure. Individual characteristics of the victims, intensity of exposure to poison and the number of animals involved, may determine slight, serious or even fatal accidents (Duarte *et al.*, 1996; Fan *et al.*, 1998; Gamborgi *et al.*, 2006; Malaque *et al.*, 2006; Riella *et al*., 2008; Basulado *et al*., 2008).

Physiological activity presented by the venom in their victims initiated several studies on its composition, in order to develop therapy for poisoning and identify potential resources for the treatment of various conditions. Among the compounds identified a protein has been well characterized, called Lopap *(Lonomia* Prothrombin Activator Protease), but the literature still lacks studies to elucidate the overall protein content of venom composition and biological fluids caterpillar, which yet remains incomplete.

Proteomes are highly dynamic and may change with development and insect environment (Wilkins *et al.*, 1997). Among the wide possibilities of bioactive molecules that insects can provide protein compounds with anti-apoptotic activity have been described (Rhee *et al.*, 2013; Choi *et al.*, 2002; Kim *et al.*, 2001; Rhee and Park, 2000). This activity may be interesting in pathologies that develop with cell loss by apoptosis, such as neurodegenerative. *L. obliqua* hemolymph anti-apoptotic activity has been described for insect cells and mammals forward physical inducers, biological and chemical (Vieira *et al.*, 2010; Mendonça *et al.*, 2008; Souza *et al.*, 2005; Maranga *et al*., 2003), but have not yet been described studies conducted with nerve cells.

This study aimed identify hemolymph and fractions protein composition of three caterpillar colonies from different cities, collected at different times and feeding on various plants, and evaluate if there were significant changes between profiles of the same. Still, the effect of *L. obliqua* caterpillars crude hemolymph and their chromatographic fractions was evaluated on cell viability of rat primary hippocampal cell cultures.

## 2. Materials and Methods

### Hemolymph collection

*Lonomia obliqua* caterpillars was collected in state of Rio Grande do Sul (South of Brazil) and frozen at −20 °C by 4 h. Hemolymph was harvested from sixty instar larvae after cutting bristles. The collected hemolymph was centrifuged by 14,000 rpm for 20 min, the supernatant was filtered through a 0.22 μm membrane filter and stored at −20 °C.

### Hemolymph Semi-Purified Fraction

Semi-purified fraction with anti-apoptotic activity was obtained from Butantan Institute (São Paulo, Brazil) (Souza *et al.*, 2005). Briefly 0.5 mL of total hemolymph was fractionated on an AKTA Purifier chromatography system equipped with a Resource Q ion exchange column (Amersham Pharmacia Biotech, USA) at a rate of 0.5 mL min^-1^ and eluted at a linear gradient (0-100%), TrisHCl 20 mmol L^-1^/Tris HCl – NaCl 1 mol L^-1^, pH 8.0. The eluate was monitored at 280, 254 and 214 nm and harvested in fractions of 1 mL.

### Animals

Wistar rats male (weighing 200 g, 100 days old) and virgin female (weighing 190 g, 90 days old) were obtained from School of Pharmaceutical Sciences/Chemistry Institute, University of São Paulo, São Paulo, Brazil. Females were caged overnight with males and copulation was verified in the morning by detection of a vaginal plug. Pregnant females were housed under controlled temperature (22±1 °C) on a 12:12 hours light/dark cycle. Food and water were provided *ad libitum*. Fetuses were obtained at 18-19 days of pregnancy (E18-E-19). Pregnant rats were anesthetized with pentobarbital (45 mg Kg^-1^) and the fetuses were rapidly euthanized by decapitation to remove their hippocampi. All experiments were conducted according to the guidelines issued by the National Council for Animal Experimentation and this study was approved by the Animal Uses Ethic Committee of the Pharmaceutical Sciences School of University of São Paulo.

### Cell culture

Rat embryonic hippocampal primary cultures were obtained according Garcia *et al.* (2012) with slightly modifications. Primary cultures were obtained by hippocampal neurons dissociation from hippocampi of E18-E19 Wistar rat embryos. Hippocampi were maintained in solution containing cooled neurobasal medium (Gibco, USA) with 100 U mL^-1^ penicillin and 100 μg mL^-1^ streptomycin. Tissues were washed with *Hank’s Balanced Salt Solution* (HBSS; Gibco, USA) and fragmented mechanically. Cell isolation was obtained by proteolytic digestion with trypsin according Jahr and Stevens (1987) and Silva *et al.* (2006). Hippocampi fragments were incubated with trypsin 0,25% (Gibco, USA), pH 7.2-7.4, at 37 °C for 10 min. Reaction was stopped with HBSS containing 277.5 U mL^-1^ DNAse (Sigma, USA) and 10% fetal bovine serum (Gibco, USA), pH 7.2-7.4. Cells were then dispersed mechanically with Pasteur pipettes of different diameters. After, hippocampal cells were resuspended in neurobasal medium (Gibco, USA) supplemented with 0.5 mmol L^-1^ L-glutamine (Gibco, USA), 25 μmol L^-1^ L-glutamic acid (Sigma, USA), 100 U mL^-1^ penicillin: 100 μg mL^-1^ streptomycin (Sigma, USA) and 2% B27 supplement (Gibco, USA), to reduce glial cell proliferation (Brewer *et al.*, 1993; Silva *et al.*, 2006). Cells suspension was then plated onto 0.01% poly-L-lysine-coated (Sigma, USA) 96-well culture plates at a density of 5.10^4^ cells per well and incubated for 7-8 days at 37°C, in humidified atmosphere of 5% CO_2_ for hippocampal neurons maturation. Half of the medium culture was replaced for the same volume of fresh medium with the same composition each 48 h. On 6-8 day, cultured cells were incubated with the treatments according to the experiment. Previous study by immunohistochemistry (Garcia *et al.*, 2012) showed a predominance of neurons in this culture, with 92% of neurons and 8% of astrocytes.

### Apoptosis induction

Hippocampal cells (5.10^4^ cells/well) were treated with 10 and 30 μmol L^-1^ H_2_O_2_ solutions [from H_2_O_2_ stock solution 30% (Merck, Germany), into supplemented neurobasal medium] fresh made, for 30 min (37°C, 5% CO_2_), after treatment with hemolymph and fractions in different concentrations. Exposure to H_2_O_2_ solutions was performed within 5 minutes of dilution, in the dark.

### Cell viability

Cells viability was evaluated by MTT-reduction assay (Mosman, 1983; Liu et al., 1997). Active metabolically cells are capable to cleave 3-(4,5-dimethylthiazol-2-yl)-2,5-diphenyltetrazolium bromide (MTT) by reduction of this yellow salt to formazan crystals, a purple compound, by mitochondrial reductases enzymes of viable cells. Formazan crystals can be measured by absorbance at 570 nm and directly correlated with viable cells number. For the assessment of cell viability, primary hipocampal cells in culture (5.10^4^ cells/well; 7-8° culture day) were incubated with hemolymph or fraction at 0.05 and 0.1% (into supplemented neurobasal medium) for 1 and 24 h (37°C, 5% CO_2_). After the period of incubation, cells were treated with 10 and 30 μmol L^-1^ H_2_O_2_ solutions (into supplemented neurobasal medium) and incubated for 30 min (37°C, 5% CO_2_). Treatments were then replaced by 100 μL 0.5 mg mL^-1^ MTT (Sigma-Aldrich Co., USA) solution and plates were incubated for 3 h (37°C, 5% CO_2_). Then, MTT solution was removed and 200 μL of dimethyl sulfoxide (DMSO, Synth, Brazil) was added to each well. After, plates were shaking for 30 min and the absorbance was measured at 570 nm in a multiwell plate reader Synergy H1 Hybrid Reader (Biotek Instruments Inc., USA).

### Statistical analysis

Data were reported as means ± SD (standard deviation) and statistical significance was evaluated by one-way analysis of variance (ANOVA) followed by the Newman-Keuls test (*p* < 0.05 was accepted as statistically significant). Experiments were conducted in quadruplicate from at least three independent experiments. Assays were analyzed by Prism 5 software (Graph Pad Software, USA).

### 2.1 Shotgun analysis

#### Sample preparation

Protein concentration of the samples was determined using a Bradford assay (Bradford, 1976) with bovine serum albumin (BSA) as a standard.

Proteins were concentrated according Wessel and Flügge (1984). Briefly, 20 μg (total proteins) of each sample were submitted to protein precipitation with chloroform and methanol (1:3:1:3, sample:methanol:chloroform:destilated water) followed by centrifugation at 17,000 rpm, 10 min. Supernatant was removed and were added three volumes of methanol (centrifugation at 17,000 rpm, 10 min). After, liquid phase was discarded and precipitated was maintained at room temperature to dry. Precipitates of concentrated proteins were resuspended in buffer urea (8 mol L^-1^), tris (0.1 mol L^-1^) and was followed the digestion protocol adapted of Klammer and MacCoss (2006). Briefly, disulfide bonds were reduced by DTT 10 mmol L^-1^ addition (37°C, 20 min) followed alkylation by IAA 50 mmol L^-1^ addition (room temperature, in the dark, 20 min). After, urea was diluted to 2 mol L^-1^ by 60 μL of tris 0.1 mol L^-1^ addition. Then, proteins were digested with trypsin (1:50, enzyme:substrate) in presence of CaCl2 1 mmol L^-1^ (37°C, 18 h). Reaction was stopped by formic acid addition (5% v/v, final concentration).

#### NanoLC LTQ-XL Orbitrap MS/MS

Chromatography separations of tryptic peptide mixture was obtained by nanoLC Ultra (nanoLC Ultra 1D plus, Eksigent, USA) equipped with autosampler nanoLC AS-2 (Eksigent, USA) and connected to LTQ-XL Orbitrap Discovery MS/MS (Thermo Fischer Scientific, USA), containing a nano-electrospray ionization source (Thermo Fischer Scientific, USA). Analytical capillary columns (100 μm × 20 cm) and pre-columns (150 μm × 2 cm) was packaged *in house* with phase-reversed C18 (5 μm ODS-AQ C18, Yamamura Chemical Lab).

Hemolymph and chromatography fractions was loaded in autosampler with injection volume 10 μL and at a flow rate of 1 μL mL^-1^ for 15 min. Step gradient mobile phase A (5% acetonitrile, 0.1% formic acid in water) was used to chromatography separations of 120 min (fractions) or 360 min (hemolymph) to mobile phase B (90% acetonitrile, 0.1% formic acid): 0-5% B in 5 min; 5-25% B in 60 min; 25-50% B in 20 min; 50-80% B in 15 min; 80% isocratic B for 5 min; 80-5% B in 1 min; 5% isocratic B for 14 min at a flow rate of 400 nL min^-1^).

Positive ion mode was employed and normalized collision energy was set to 35%. Full scan MS spectra were acquired from *m/z* 400-1,600 at a resolving power of 30,000, followed by acquisition of 8 MS2 mode spectra of the most abundant ions. Fragmentation was obtained by dissociated induced collision (CID), with *Q* activation = 0.250, time of activation = 30 ms and isolation amplitude = 1 Da. Acquired peptide masses were added to dynamic exclusion list with size of 100 ions and time of permanence of 30 s during acquisition of MS2 spectra. Spray voltage used were 2.2 kV, capillary temperature of 275°C, capillary voltage of 34 V, and the collision gas used was helium.

#### Data analysis

For protein identification, a no-redundant sequences data bank referred to entries of individuals proteins for *“Lonomia”* presents in NCBI data bank (http://www.ncbi.nlm.nih.gov) was constructed. From this bank, RAW files of MS-2 spectra was searched using *Comet* search software (Eng *et al.*, 2013). Identification proteins and peptides was obtained using *Pattern Lab for Proteomics* software from homology with data bank sequences (Carvalho *et al.*, 2012). Candidates peptides were considered those containing one or two tryptic ends. Cysteine carbamidomethylation was adopted as fixed modification. Tolerance of 50 ppm to precursors ions and 1 Da to fragment ions were employed on the data search.

#### Search Engine Processor

software (Carvalho *et al.*, 2012) was used to filter identified spectra. Parameters *Xcorr*, *DeltaCN*, *DeltaMass*, *Peaks Matched* e *Spec Count Score* were used to generate a Bayesian score. Cutoff was established to accept 1% false-positive, based on reversed data base identifications. Furthermore, 6 residues were adopted as minimum sequence length. Results were post processed to set identifications less than 10 ppm of mass variation.

## 3. Results and discussion

### 3.1 Hemolymph and chromatographic fraction protein identification by NanoLC MS/MS

Studies were conducted using a gel-free proteomic strategic in a nanoLC/LTQ-Orbitrap system. A total of 71 distinct hemolymph proteins were identified from 3 lots of caterpillars, with 34 common proteins between the same. In fractions was found 40 total distinct proteins, with 9 shared between the 3 lots. A Venn diagram displays the results in **Fig. 1**.

**Fig. 1.**
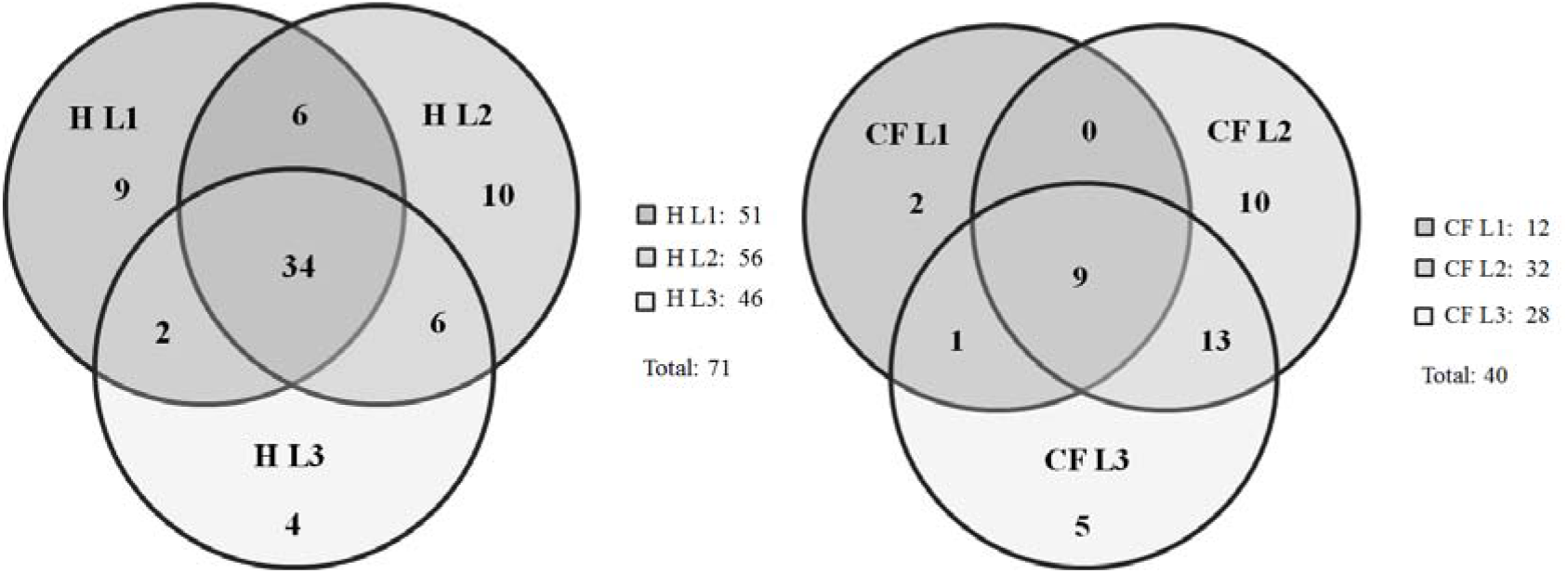
Venn diagram depicting the total number of proteins identified in 3 lots of hemolymph (H) and chromatographic fractions (CF) as well as the number of overlapped proteins.

**Fig. 2.**
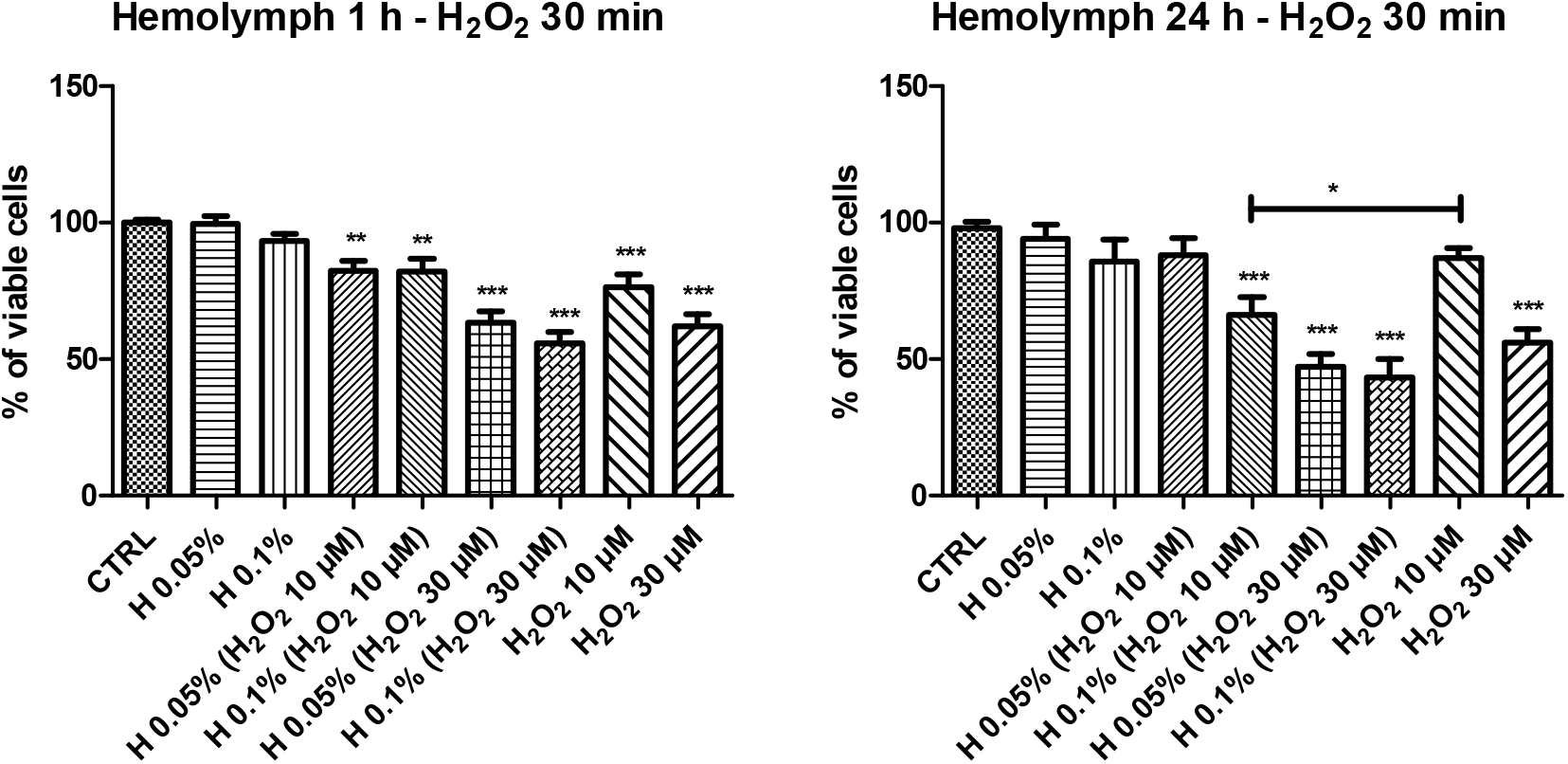
Evaluation of cell viability by MTT assay. CTRL (Control; n = 5 wells per group in each of the three independents assays); H (Hemolymph; concentrations at 0.05% or 0.1% v/v); H_2_O_2_ (Positive control; concentrations at 10 or 30 μM). \**p* < 0.05, \*\**p* < 0.01 and \*\*\**p* < 0.001 compared with CTRL and intergroup comparison (ANOVA and Newman-Keuls multiple comparison).

Considering hemolymph and CF samples, 76 different proteins were identified, with molecular weights ranging from 2,107.1 to 71,540.4 Da: 71 in hemolymph and 40 in CF, with 34 common proteins between then. Five proteins were found only in CF and 36 just in hemolymph.

Related to CF, a proportionately larger number of peptides was generated by hemolymph, due to the larger number of proteins present in this. The number of peptides produced by proteolytic digestion is proportional to abundance of its protein in a sample (Liu *et al.*, 2004). Thus, peptides in larger quantity were selected to MS2 more frequently, generating a larger number of spectral counts which is proportional to the abundance of each protein in the sample.

Five proteins were identified only in CF, since in the crude sample, more peptides can compete to the selection of fragmentation within the same cycle of acquisition, and thus, peptides signal with less intensity can be suppressed at the expense of others compromising their identification.

Protein identification in *L. obliqua* samples by shotgun analysis is impaired by the limited range of data contained in the database for this specie. Proteins amino acids sequences data found in databases are mainly from transcriptomic studies (Veiga *et al.*, 2005).

Serine proteases (45%), hemolins (38%), protease inhibitors (7%) and antiviral protein (5%) were found in CF, as well as sensory proteins, heat shock proteins, cysteine proteases and lectins less expressively.

In hemolymph were found predominantly antiviral protein (24%), serine proteases (23%), hemolins (16%) and protease inhibitors (10%). Heat shock proteins, serpins, sensory proteins, oxidoreductases, lectins, lyases and ribonucleases also were identified. Other found proteins together represent 20% of the total, and are structural proteins or proteins involved in metabolic functions.

Significant qualitative and quantitative variations in hemolymph amino acids concentration and protein composition were related to insect age and stage of development (Roman and Jegorov, 1991; Whitmore and Gilbert, 1974), diet (Ortel, 1995) and ambient temperature (Roman and Jegorov, 1991; Cui *et al.*, 2011). Thus, this can have determined differences in protein composition observed in the three lots of hemolymph.

**Table 1.**
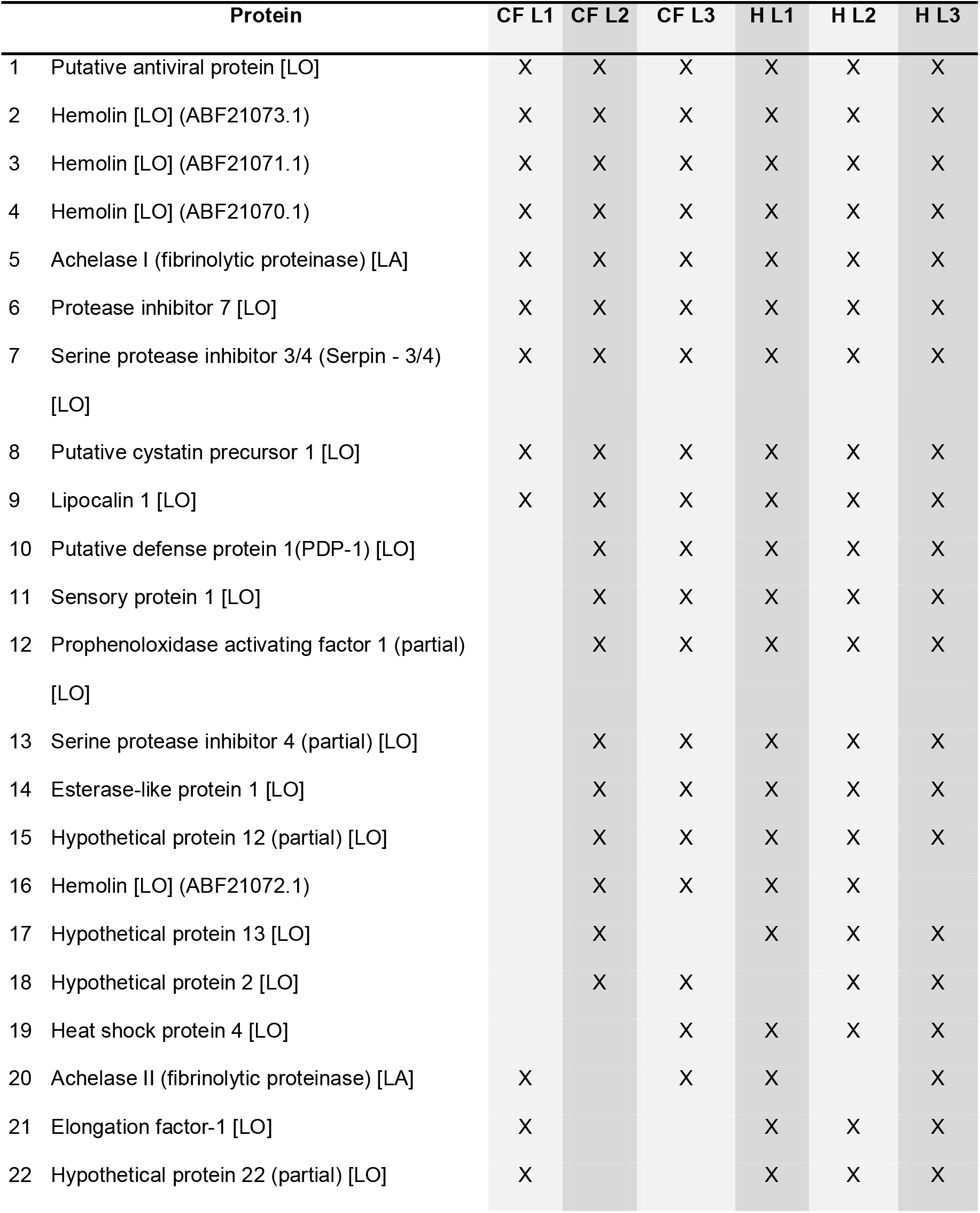
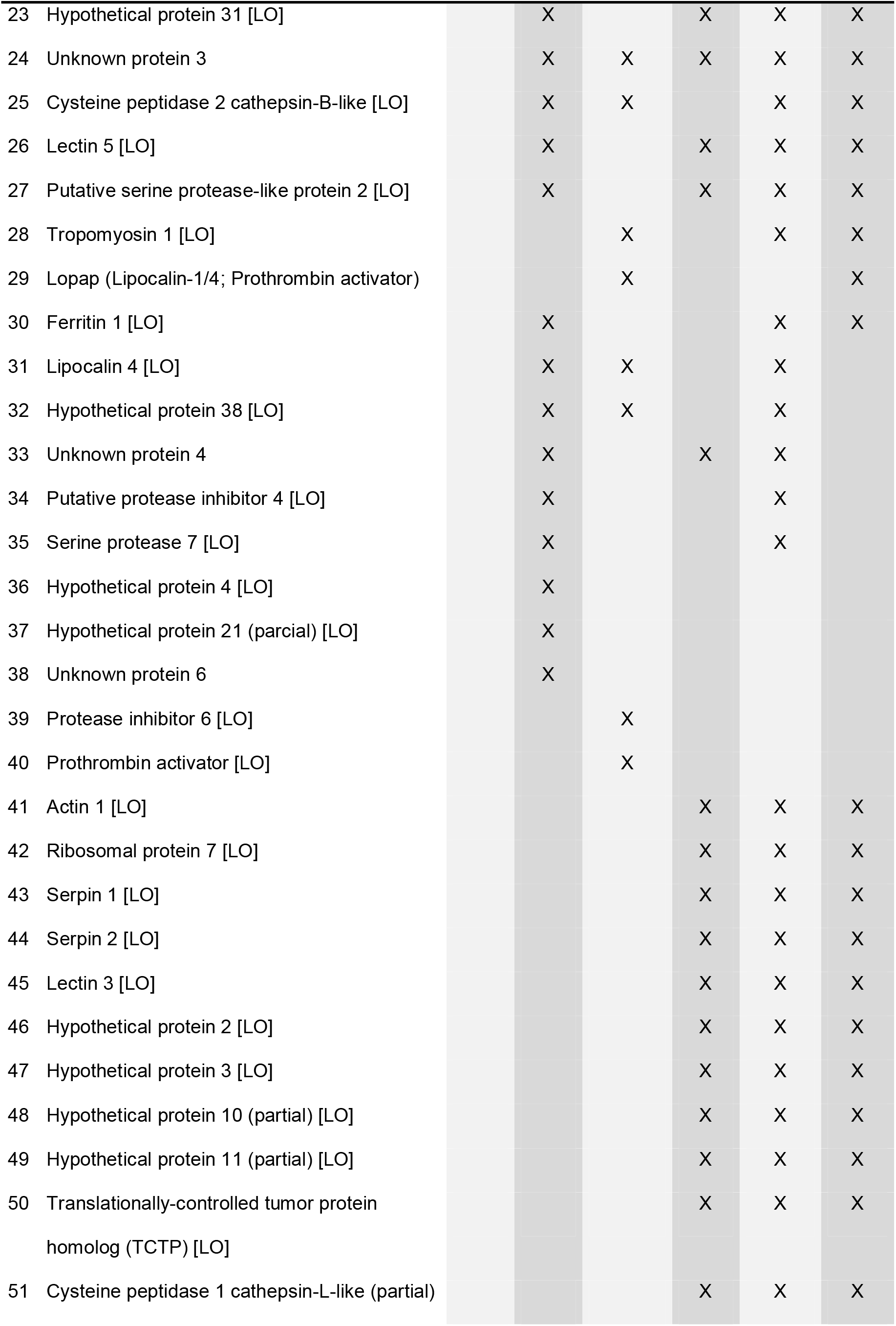
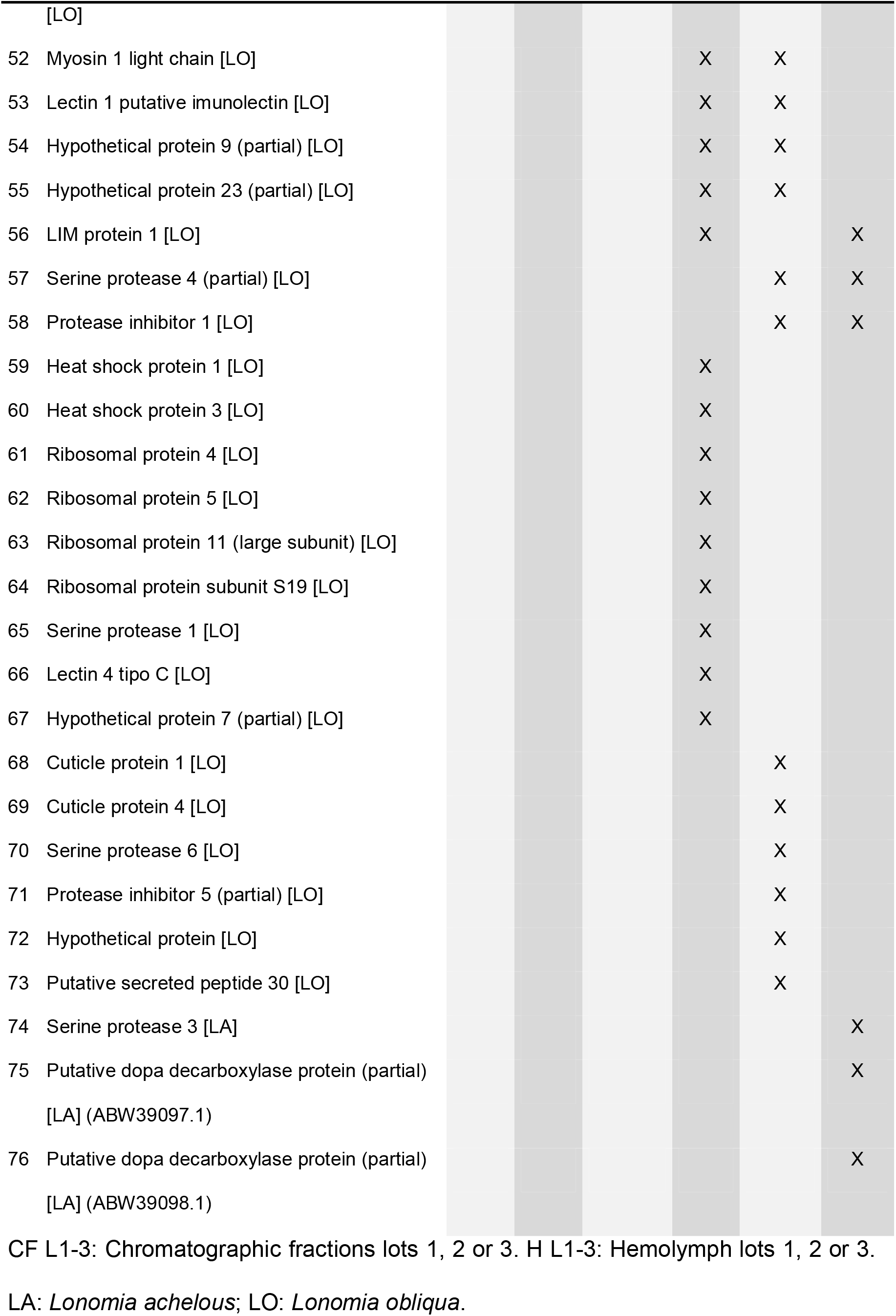
Identified proteins on hemolymph and chromatographic fractions samples by *shotgun*.

Insects present a metamorphosis process where a combination of grow sinaling, activation, differentiation and specific tissues physiological apoptosis is highly regulated. Probably substances responsible for this signaling circulate in hemolymph, which makes their interesting for study to identification of bioactive compounds with biotechnological potential (Maranga *et al.*, 2003).

### 3.2 Hemolymph and CF effects on H_2_O_2_-treated cells

#### 3.2.1 Hemolymph

Mitochondrial metabolism of 0.05% and 0.1% hemolymph treated cells for 1 h and subsequently subjected to apoptosis induction with H_2_O_2_ 10 μmol L^-1^ was significantly reduced (*p* < 0.01) compared to control without inducing apoptosis. However, when compared to apoptosis induction control with the same H_2_O_2_ concentration, there was no significant difference. Viability of 0.05% and 0.1% hemolymph cells treated for 1 h with apoptosis induced by H_2_O_2_ 30 μmol L^-1^ also had a significant reduction (*p* < 0.001) relative to untreated control, but there was no significant difference when the comparison was made with H_2_O_2_ 30 μmol L^-1^ control.

Treatment with 0.05% hemolymph, H_2_O_2_ 10 μmol L^-1^ showed no statistical difference compared to control and H_2_O_2_ 10 μmol L^-1^. Cells submitted to 0.1% hemolymph, H_2_O_2_ 10 μmol L^-1^ had significant reduction (*p* < 0.001) in cell viability compared to control and H_2_O_2_ 10 μmol L^-1^ (*p* < 0.05).

Cells with 0.05% and 0.1% hemolymph for 24 h, exposed to H_2_O_2_ 30 μmol L^-1^ had significantly reduction viability (*p* < 0.001) relative to control and was not difference in comparison with H_2_O_2_ 30 μmol L^-1^.

Presence of other active substances that may play an inhibitory effect on cell viability and proliferation, such as A2 phospholipase involved in hemolytic activity presented by the poison and hyaluronidase activity should be considered in evaluating the crude hemolymph effect (Heinen *et al.*, 2014).

#### 3.2.2 Chromatographic Fraction (CF)

Treated cells with 0.05% and 0.1% CF for 1 h and H_2_O_2_ 10 μmol L^-1^ produced viable cells significant reduction (*p* < 0.001 and *p* < 0.01 respectively) compared to untreated control and 0.05% and 0.1% CF for 1 h, H_2_O_2_ 30 μmol L^-1^ (*p* < 0.001). There were no significant differences when comparing CF and H_2_O_2_ 10 μmol L^-1^, either between CF and H_2_O_2_ 30 μmol L^-1^.

For 0.05% and 0.1% CF (24 h), H_2_O_2_ 10 μmol L^-1^ there is not statistical difference compared to control, however occurred cell viability increase related to H_2_O_2_ 10 μmol L^-1^ (*p* < 0.05 and *p* < 0.01, respectively). Cells with 0.05% and 0.1% for 24 h, H_2_O_2_ 30 μmol L^-1^ had significant reduction viability (*p* < 0.01) regarding control.

Increase in viable cells (*p* < 0.05) was produced by 0.05% CF, H_2_O_2_ 30 μmol L^-1^ related to H_2_O_2_ 30 μmol L^-1^ while 0.1% CF, H_2_O_2_ 30 μmol L^-1^ had no difference.

**Fig. 3.**
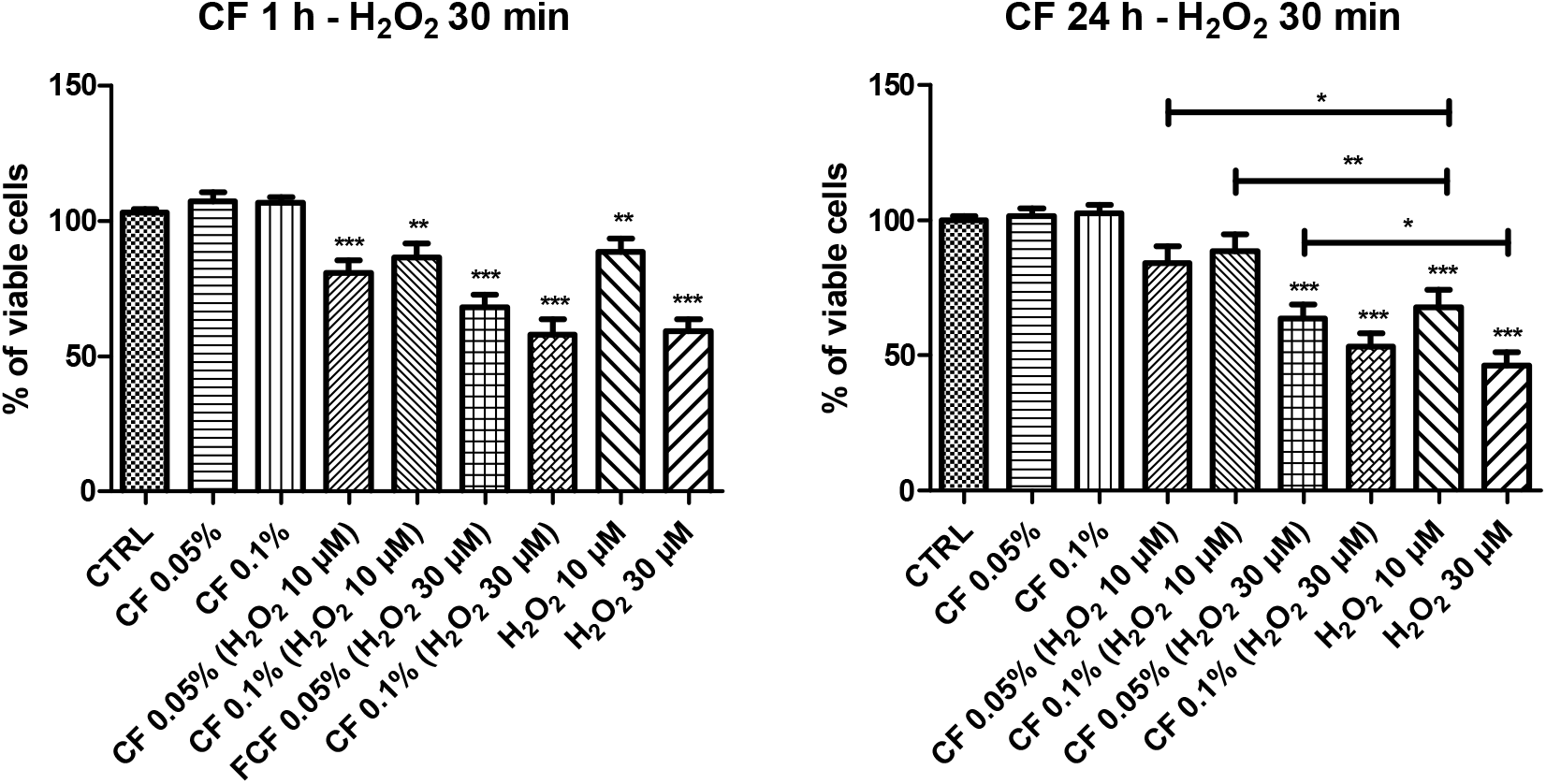
Evaluation of celL viability by MTT assay. CTRL (Control; n = 5 wells per group in each of the five independents assays); CF (Chromatographic fraction; concentrations at 0.05% or 0.1% v/v); H_2_O_2_ (Positive control; concentrations at 10 or 30 μM). \**p* < 0.05, \*\**p* < 0.01 and \*\*\**p* < 0.001 compared with CTRL and intergroup comparison (ANOVA and Newman-Keuls multiple comparison).

After 24 h of exposure the rat hipocampal neurons culture to CF, and subsequently exposure to H_2_O_2_ for 30 min, the cellular viability kept higher than positive control (only H_2_O_2_), but a reduction in neuronal viability was observed related to control (without oxidative induction).

Several studies have shown the property of substances from insects hemolymph in the inhibition of apoptosis promoted by different inducers (physical, chemical and biological), in insects and mammals cell cultures (Rhee *et al.*, 2013; Vieira *et al.*, 2010; Mendonça *et al.*, 2008; Souza *et al.*, 2005; Maranga *et al.*, 2003; Choi *et al.*, 2002; Kim *et al.*, 2001; Rhee and Park, 2000). The effect in a variety of cells opposite the apoptosis induction by multiple mechanisms suggests that it may act in some apoptosis conserved way (Heinen *et al.*, 2014).

*L. obliqua* hemolymph anti-apoptotic activity was observed in Sf-9 insect cells (Souza *et al.*, 2005; Maranga *et al.*, 2003; Vieira *et al.*, 2010), in HEK-293 human cells (Mendonça *et al.*, 2007) and in mammalian cells V-79 (Heinen *et al.*, 2014). Mendonça *et al*. (2007) observed that exposure to hemolymph and chromatographic fractions produced high electrochemical potential in the mitochondrial membrane maintenance.

Cytoprotective activity was described too in leucocytes and endothelial cells (Chudzinski-Tavassi *et al.*, 2010), in HUVECs (Fritzen *et al.*, 2005) and in fibroblasts FN-1 *(rLosac –* recombinant *Lonomia obliqua* Stuart-Factor Activator) (Alvarez-Flores *et al.*, 2012), but associated to different proteins from *L. obliqua*.

Further studies to verify expressed proteins from samples treated cultures may indicate if factors such as cells from animals tissues adaptation differences to *in vitro* condition can be related to results obtained. Moreover, it is still unknown anti-apoptotic protein interaction with specific cellular receptors that may not be present or not be available in quantities sufficient for the manifestation of the effects in the studied cell. The structure of the anti-apoptotic effects responsible protein remains without being elucidated and thus the prediction of their susceptibility to degradation or inactivation are also compromised.

## 4. Conclusion

As demonstrated in this study, animal venoms protein composition such as *L. obliqua* can provide qualitative and quantitative variations intra species when compared different origins animals and consequently exposed to various environmental factors. Concentration of free ions naturally present in the hemolymph can likewise differ between animals and produce shift in the elution of specific proteins during ion exchange chromatography making not reproducible elution pattern of the fractions. Thus, proteins present in fraction used in this study may differ over those described in the literature using the same fractionation strategy. Still, there may be synergistic action between the substances present in the hemolymph, which may contribute to a greater or lesser effect.

This study describes too, for the first time, the evaluation of *L. obliqua* hemolymph and fraction on cell viability of neuronal cells primary culture. The results show that treatment of primary embryonic cultures of Wistar rats hipocampal neurons with CF for 24 h and subsequently exposure to H_2_O_2_ for 30 min, showed increased cell viability compared to the positive control group, treated only with the inducing agent H_2_O_2_ but they have significantly reduced mitochondrial metabolism when compared to the control without oxidative induction. The other conditions and treatment tested concentrations did not produce evidence that the hemolymph and fractions have produced a neuroprotective effect.

## Acknowledgments

The autors aknowledge the financial support by Brazilian agency CAPES. The autors gratefully aknowlege the support of Dr. Lisete M. Lorini, from University of Passo Fundo, Passo Fundo, Brazil, the Rio Grande do Sul Toxicology Information Centre, Porto Alegre, Brazil, Dr. Mariana R. Ely and Dr. Edegar Fronza, from University of Caxias do Sul, Caxias do Sul, Brazil.

